# Altered auditory feedback induces coupled changes in formant frequencies during speech production

**DOI:** 10.1101/733121

**Authors:** Ding-lan Tang, Daniel R. Lametti, Kate E. Watkins

## Abstract

Speaking is one of the most complicated motor behaviours, involving a large number of articulatory muscles which can move independently to command precise changes in speech acoustics. Here, we used real-time manipulations of speech feedback to test whether the acoustics of speech production (e.g. the formants) reflect independently controlled articulatory movements or combinations of movements. During repetitive productions of “head, bed, dead”, either the first (F1) or the second formant (F2) of vowels was shifted and fed back to participants. We then examined whether changes in production in response to these alterations occurred for only the perturbed formant or both formants. In Experiment 1, our results showed that participants who received increased F1 feedback significantly decreased their F1 productions in compensation, but also significantly increased the frequency of their F2 productions. The combined F1-F2 change moved the utterances closer to a known pattern of speech production (i.e. the vowel category “hid, bid, did”). In Experiment 2, we further showed that a downshift in frequency of F2 feedback also induced significant compensatory changes in both the perturbed (F2) and the unperturbed formant (F1) that were in opposite directions. Taken together, the results demonstrate that a shift in auditory feedback of a single formant drives combined changes in related formants. The results suggest that, although formants can be controlled independently, the speech motor system may favour a strategy in which changes in formant production are coupled to maintain speech production within specific regions of the vowel space corresponding to existing speech-sound categories.

**New & Noteworthy:** Findings from previous studies examining responses to altered auditory feedback are inconsistent with respect to the changes speakers make to their production. Speakers can compensate by specifically altering their production to offset the acoustic error in feedback. Alternatively, they may compensate by changing their speech production more globally to produce a speech sound closer to an existing category in their repertoire. Our study shows support for the latter strategy.

## Introduction

Speaking is one of the most complicated motor behaviours. Fluent speech production requires fine coordination of many muscles and integration with simultaneous auditory and somatosensory feedback. How this coordination of movement is achieved given the large number of muscles involved and degrees of freedom of movement and how these map on to sensory targets are largely unresolved problems in the field of speech motor control. On the sensory side, it is clear that auditory feedback is particularly important in speech development; pre-lingually deaf children have difficulties with speech articulation (Svirsky et al., 2004). It is also important for the maintenance of speech intelligibility, which can deteriorate in post-lingually deaf adults (Waldstein, 1990). Such individuals can rely on somatosensory feedback from orofacial muscles to achieve normal speech articulation (Nasir and Ostry 2008). Somatosensory feedback is also relied on by some individuals with normal hearing to monitor their speech production (Lametti et al., 2012; Tremblay et al., 2003).

The relationship between the movements used to articulate speech sounds and the sensory consequences is learned during speech development and used to build internal models that encode these sensorimotor mappings and guide speech production. Internal models allow the brain to predict the sensory consequences of actions, perform online correction and rapidly detect errors. The exact nature of these representations is unknown. That is, are speech sound targets (e.g., phonemic categories) represented by a combination of movements used to achieve the sensory goal—or are the targets of speech the consequence of a series of independently controlled articulatory movements?

Internal models need to be updated and trained to adapt for changes in the environment or in our bodies that affect speech perception and production—for example, when imitating the accent of an interlocutor or coping with the changes in shape of the vocal tract during adolescence. It is possible to explore these models experimentally by manipulating sensory feedback in real-time and measuring the ways in which speakers adapt their speech production (Houde & Jordan 1998; MacDonald et al., 2011; Lametti et al., 2018a).

The basic speech adaptation paradigm involves altering frequencies that characterize vowel sounds, feeding these back to the speaker in near real time (e.g., with less than a 40 ms delay), and measuring the compensatory changes in the vowel sounds the speaker subsequently produces (Houde and Jordan 1998, 2002; Purcell and Munhall, 2006a, b; Mollaei et al., 2013). In speech, vowel sounds are defined by vocal tract resonances known as formants. Formants are distinctive frequency components evident in the acoustic signal reflecting the size and shape of different chambers in the vocal tract. The first two formants (F1 and F2) in a vowel sound contain most of the acoustical energy and are sufficient to disambiguate vowel sounds. By examining the relationship between the changes in the first two formants of vowels produced when speech feedback is altered, we can gain insight into how speech movements are controlled. For example, if the frequency of F1 for the vowel in “head” is increased, it sounds closer to the F1 frequency of the vowel in “had”. Participants can compensate by lowering the frequency of F1 in their subsequent productions, which would counteract the acoustic change applied and result in an F1 that with the frequency shift applied and fed back sounds closer to that of the intended vowel in “head”. Alternatively, participants could change the frequency of both F1 and F2 in their productions; a combination of decreasing F1 frequency whilst simultaneously increasing F2 could move their production closer to that of the vowel in “hid”, a familiar region of the vowel space (see Figure 1). Determining whether adaptive speech movements are characterized by specific changes along separate dimensions, (e.g., F1, which corresponds to tongue height, and F2, which corresponds to front-back position of the tongue body; Ladefoged, 2001) or are controlled more globally by the production of vocal tract shapes that involve both dimensions, would inform our understanding of the nature of representations in models of speech motor control.

**Figure 1.**
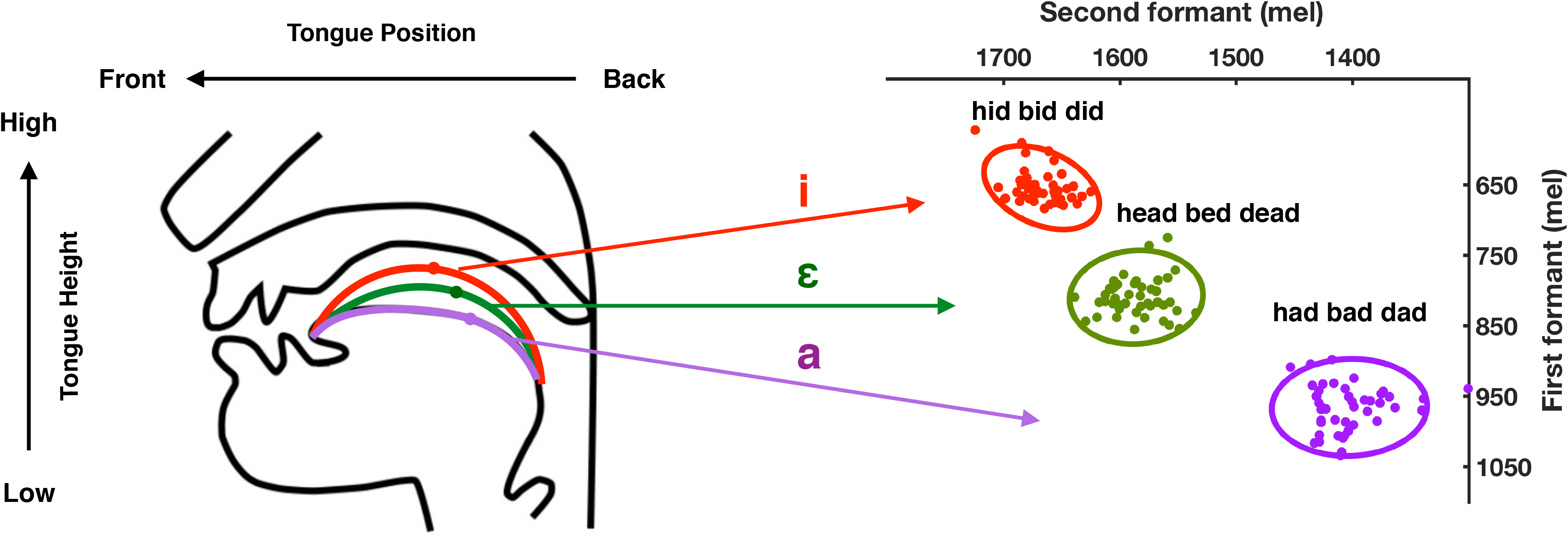
Vowel production. Schematic showing the relationship between tongue height and position in the vocal tract (left) and the corresponding frequencies of the first and second formants of vowels (/I/, /ɛ/, and /æ/) in vowel space from a single participant in the current study (right). Data points correspond to F1 and F2 values of single utterances during the vowel exploration phase in a representative participant. The ellipses represent a 90% confidence interval around the data points of same colour.

The current literature provides evidence for both specific changes in F1 during adaptation to F1 alterations and more global strategies involving changes in both F1 and F2. Previous work involving adaptation to gradual shifts in F1 showed compensation in F1 production but no significant change in F2 (Villacorta et al. 2007). In contrast, another study found changes in both F1 and F2 (MacDonald et al., 2011) when feedback in F1 was changed abruptly. In this case, however, the direction of change in F2 was the same for both upwards and downwards shifts of F1 and occurred more gradually than the change in F1. This suggests that F1 compensation, which is directional and compensatory for the applied F1 shift, is independent of the F2 change. On the other hand, there is also evidence to suggest that formants are not modified independently. For example, reliable F2 changes in response to shifts in F1 were observed in the opposite direction to the compensatory changes in F1 for utterances containing the vowel /⍰/, such as “bed” and “head” (Rochet-Capellan and Ostry, 2011; Lametti et al., 2018b).

The inconsistency in results of previous studies could be due to the relatively small sample sizes and the large variability among participants. A sample size of 12 (or less) per group is typical for these kinds of studies yet a large proportion of people (10-40%) exhibit little or no compensation (Lametti et al., 2012, 2014; Rochet-Capellan et al. 2012). In addition, meta-analyses of such studies indicate that on average 22% of participants altered production in a pattern that follows (instead of opposes) the direction of the F1 shift (MacDonald et al., 2011). These individual differences could be explained by preferences for reliance on auditory or somatosensory feedback during speech perturbation studies (Lametti et al., 2012) and further complicates work aiming to explore the relationship between F1 and F2 production in response to alterations of a single formant (MacDonald et al., 2011).

In the current study, we revisit the idea that speakers counteract errors in formant production by only changing the altered formant. In our first experiment, we increased the frequency of F1 and examined compensatory changes in both F1 and F2. Based on previous findings (see above), we predicted that participants would decrease their F1 to offset the perceived acoustical error. We also predicted based on studies where the shift was applied abruptly (e.g. Rochet-Capellan and Ostry 2011; Lametti et al., 2018b) that increases in F2 would accompany the F1 decrease to move speech closer to a known region of the vowel space. In other words, when participants produced “head, bed, dead” and heard an upward shift in F1, an increase in F2 concurrent with the compensatory decrease in F1 would place the new production closer to “hid, bid, did” in vowel space (see Figure 1). In a second experiment, we decreased the frequency of F2 and examined compensatory changes in both F2 and F1. We predicted that increases in F2 would be observed to offset the induced acoustical error, and, as for responses to increased F1, decreases in the unaltered formant (this time F1) would accompany these F2 increases to move speech closer to a known region of the vowel space (Figure 1). Such a result would support the idea that, although F1 and F2 can be controlled independently (MacDonald et al., 2011), the speech motor system favours a strategy that moves formant production to specific regions of the vowel space.

## Methods

### Participants and Apparatus

Forty native English speakers between the ages of 18 and 40 years, with no reported history of speech, language, or hearing disorders took part in the study. The Central University Research Ethics Committee (at the University of Oxford) approved the experimental protocol and participants gave informed consent. There were twenty participants in Experiment 1 (21.55 ± 5.27 years; 18 females) and 20 different participants in Experiment 2 (20.15 ± 3.59 years, 14 females).

### Apparatus

The experiment was conducted in a quiet room with participants seated in front of a computer screen, which was used to present words. During the experiment, participants wore over-ear headphones (Sennheiser, HD 280 Pro) and a head-mounted microphone (Shure, WH20) that was placed approximately 4-7 cm away from the right corner of the participant’s mouth. A Matlab Mex-based program Audapter (Cai et al., 2008; Tourville et al., 2013) was used to alter speech and play it back to participants with an unnoticeable delay.

### Procedure

In both experiments, consonant-vowel-consonant words were presented on the computer screen for 1500 ms, one at a time. The inter-trial-interval was 750 ms. Participants were told that they would have to read out the words on the screen, and that they would hear their own voice through headphones.

First, participants completed a vowel exploration session with normal feedback in which they produced nine different words (‘dead’ ‘bed’ ‘head’, ‘dad’ ‘bad’ ‘had’, ‘did’ ‘bid’ ‘hid’, containing three different vowels /⍰/, /æ/ and /I/, correspondingly) 15 times each. This session allowed participants to explore the vowel space. It also provided us with estimates of F1 and F2 frequencies for each vowel category in each individual. Vowel exploration was followed by three experimental phases: Baseline, Learning, and Unlearning (Figure 2). During the baseline phase, three different words “dead, bed, head”, containing the same vowel /⍰/, were produced 15 times each by participants with normal feedback (2 min). In the learning phase (10 min), participants produced “dead, bed, head” 75 times each with altered auditory feedback designed to induce sensorimotor learning (see Real-Time Feedback Alteration). In the final phase, Unlearning, “dead, bed, head” were produced 30 times each with normal feedback (4 min). The order of words was randomized within each phase.

**Figure 2.**
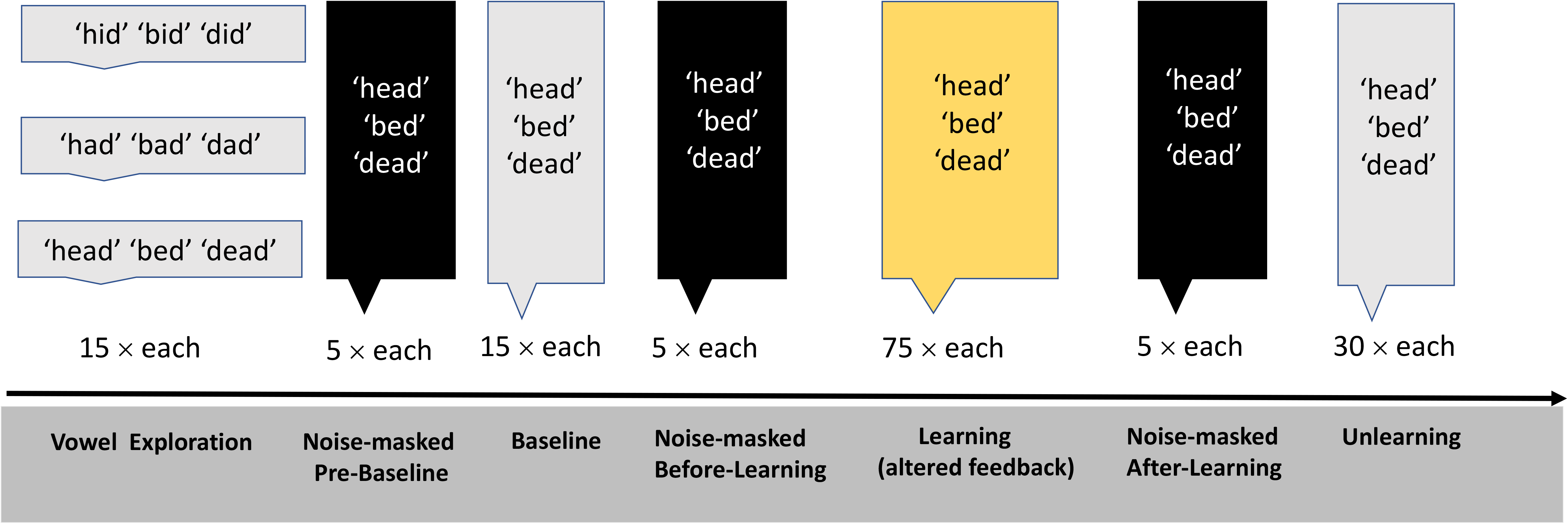
Experimental procedure. The study involved Vowel Exploration, Baseline, Learning, and Unlearning, interleaved with three short noise-masked sessions.

Three short noise-masked sessions preceded each of these experimental phases. During these noise-masked sessions, participants produced the words “dead, bed, head” five times each but, instead of hearing feedback of their speech, they heard masking noise that scaled with the amplitude of the produced speech signal (i.e. the signal-to-noise ratio was 0 dB). White noise was also added to block participants’ ability to hear their own speech as much as possible.

The three noise-masked sessions were carried out at the following time-points: 1) Pre-Baseline 2) Before-Learning, and 3) After-Learning. The aim of these sessions was to measure whether feedforward changes in speech production associated with sensorimotor learning would persist in the absence of auditory feedback. Figure 2 shows a schematic of the Experimental Procedure.

### Real-Time Feedback Alteration

To induce sensorimotor learning in speech, the frequency of either F1 or F2 was altered and played back to participants with an imperceptible delay. The formant alteration was applied in mels — a perceptual scale characterizing changes in frequency that are judged by listeners to correspond to equal changes in pitch (Stevens et al., 1937). The transformation from Hz to mel used was:

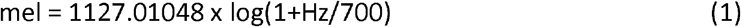

Participants were randomly assigned to one of two groups. The F1+110 group (Experiment 1) experienced a 110 mel increase in first formant (F1) production, while the F2−110 group (Experiment 2) experienced a 110 mel decrease in second formant (F2) production. A feedback shift magnitude of 110 mel was selected for this study because it is comparable to the F1 feedback perturbation used previously (Lametti et al., 2018b).

Both the increase in F1 frequency (F1+110) and the decrease in F2 frequency (F2−110) of the vowel /⍰/ in the produced words resulted in the percept of speech with formant frequencies that were closer to F1 or F2 frequencies respectively of the vowel /æ/ in the vowel space (Fig. 1). Compensatory responses were expected to be produced by speakers to counteract the perceived shift, either by reducing F1 (Experiment 1: F1+110 group) or increasing F2 (Experiment 2: F2−110 group) (Fig. 1). We were also interested in the extent to which speakers altered their production of the unshifted formant in each experiment.

### Auditory analysis

Speech was recorded at 16000 Hz in MATLAB and analysed using custom-written MATLAB scripts. First, the F1 and F2 frequencies of each vowel were extracted from the data output of Audapter, which estimates the first two formant frequencies using linear predictive coding. Then the average formant frequency was calculated from a 30-ms segment at the centre of each vowel. Utterances that contained production errors or contained formant frequencies that were greater than three standard deviations from a participant’s mean F1 and F2 values in each phase of the experiment were excluded from further analyses. This resulted in approximately 3% of the data being excluded.

### Statistical analysis

In each participant, the F1 and F2 frequency data for utterances produced during the learning and unlearning phases of the experiments were normalised by subtracting the mean and dividing by the standard deviation of the formant frequencies produced during baseline (with normal feedback) to create a Z value for each datapoint.

Following Lametti et al. (2018b), we assessed the significance of compensatory changes in the shifted and unshifted formant frequencies in each experiment using a one-sample t-test on the average Z values of the last 30 trials in the learning phase (against z = 0, no change from baseline). As the directions of the compensatory change to the formant feedback shifts were predictable, we used directional t-test with a significance level of p < .05. In the current study, we also used noise-masked sessions before the baseline, before and after the learning phase. Data obtained in these sessions allowed us to compare the effect of the perturbation on speech production under the same conditions when feedback was blocked. The pre-baseline session was used to normalise the before- and after-learning sessions as described above (we first confirmed that there was no change in F1 and F2 in the sessions performed either side of the baseline; p > .05). Then average change from before-learning session in produced F1 and F2 during after-learning session was calculated and compared with zero using directional t-tests to examine if there was a significant change from before to after-learning.

Following MacDonald et al. (2011), we assessed the significance of the return-to-baseline changes in the shifted and unshifted formant frequencies in the unlearning phase of each experiment. The average Z values of the last 15 trials in the unlearning phase were compared using a one-sample t-test (against Z = 0, i.e., the baseline average).

For each participant, we also calculated the Euclidean distance between the centre of the vowel clouds for “hid, bid, did” produced during the vowel exploration phase of the experiment and for “head, bed, dead” produced at baseline and at the end of (last 30 utterances) the learning phase (Lametti et al., 2018b). Learning-related changes in this distance were then compared to zero (no change) with one-sample t-tests to examine whether there was a significant change in the distance induced by F1 (experiment 1) or F2 (experiment 2) feedback perturbation. Distance changes were also compared between groups (experiment 1 and experiment 2) using a two-tailed t-test.

### Unplanned Analyses

In an exploratory analysis, we compared the direction of compensation in Experiment 1 and Experiment 2. To do this, we calculated the angle between the vector that represents the direction of actual compensatory change from baseline and the vector representing the direction to move from “head, bed, dead” to “hid, bid, did” for each participant. The angles were then compared to zero (no difference from required angle) with one-sample t-tests to examine whether there was a significant difference between actual movement direction and the direction required in each Experiment separately. The angles were also compared between groups (experiment 1 and experiment 2) using a two-tailed t-test.

## Results

### Experiment 1: F1 shift of +110 mel

Figure 3A shows how F1 and F2 produced by participants, normalized as z-scores, changed during the sensorimotor learning and unlearning phases in response to a 110 mel increase in F1. By the end of the learning phase, participants had decreased their F1 on average by 30 mel to compensate for the increase in F1 and increased F2 on average by 15 mel (Figure 3B). Directional t-tests showed that both F1 and F2 frequency at the end of the learning phase (last 30 utterances of learning phase) were significantly different from zero (baseline average) (F1: t(19) = 6.24, p < .001, d = 1.97; F2: t(19) = 2.88 p = .005, d = 0.91).

**Figure 3.**
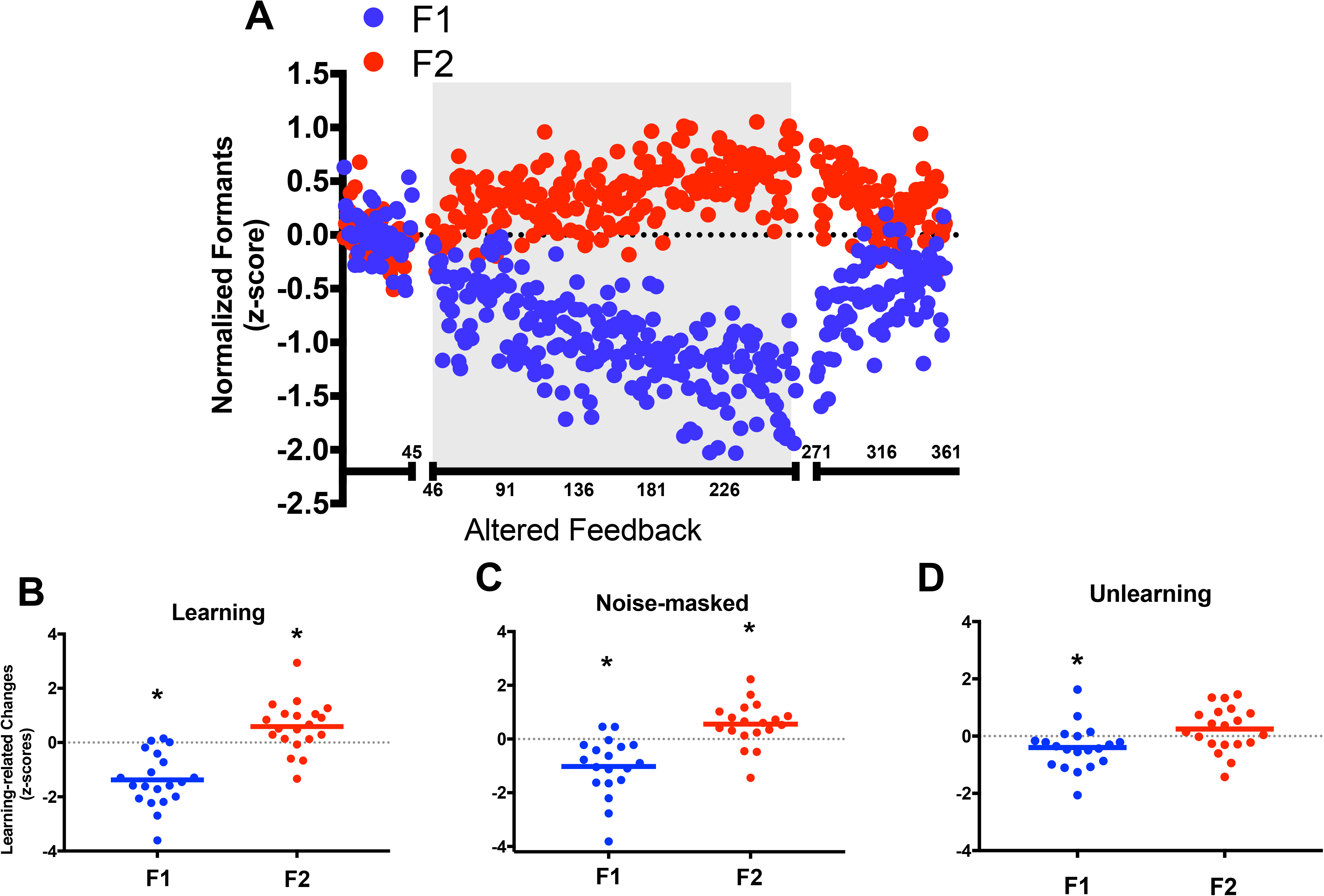
Experiment 1: F1 shifted upwards by 110 mel. Change in normalized formant frequencies during speech production. Values for F1 are shown in blue and F2 in red. The dashed line reflects no change from baseline. The graph in the top panel (A) shows the group average changes across learning and unlearning phases of the experiment relative to baseline; solid circles indicate the group average for each trial; the shaded area represents the trials where the altered feedback was applied. The graphs in the bottom panel (B, C and D) show average changes in normalised formant frequencies for individual participants for the Learning (B), Noise-masked (C) and Unlearning (D) phases of the experiment relative to baseline; the solid lines represent the group mean; * indicates significant change from baseline.

Compensatory responses of F1 and F2 production were also evaluated under noise-masked conditions, which were used as a measure of changes from before to after learning in the same context (i.e. with no feedback). Figure 3C shows the change in normalized F1 and F2 for heavily noise-masked speech from before and after learning. After learning, F1 significantly decreased (t(18) = 4.12, p < .001, d = 0.98) and F2 significantly increased (t(18) = 3.00, p = .004, d = 0.69).

By the end of unlearning phase, F2 production returned to baseline (one-sample t-test against baseline average, Z = 0: t (19) = 1.42, p= .172), while F1 remained significantly reduced relative to baseline (t (19) = 2.33, p= .031) after these 90 trials with normal feedback (Figure 3D).

In sum, in response to an F1 feedback perturbation, both F1 and F2 in production were observed to change though the magnitude of the change in the shifted formant (F1) was larger than in the unshifted one (F2). The results provide support for the idea that a single formant perturbation will induce a combination of changes in the shifted formant as well as the unshifted formant. Such combined changes could place the new production closer to a learnt pattern of speech production used to produce another vowel sound (“hid, bid, did” in current study; Figure 5A). This hypothesis was tested by comparing the magnitude (Euclidean distance) and direction of the compensatory shifts in F1 and F2 in vowel space between the pre- and post-learning productions of (“head, bed, dead”) and the production of “hid, bid, did” during the vowel exploration phase of the experiment before learning had occurred.

As shown in Figure 5B, participants in Experiment 1, responding to an increase in F1 feedback, moved their productions of “head, bed, dead” closer to the vowel space occupied by “hid, bid, did” by a magnitude of 32 mel on average (one-sample t-test against zero, t(19) = 5.44, p < .001, d =1.72). The direction of this change was not significantly different from that predicted (Figure 5C; t(19) = 1.45, p = .164). The same results were obtained by analysis of the data from the noise-masked session after learning.

### Experiment 2: F2 shift of −110 mels

In Experiment 2, we gave participants feedback with a downshift in F2 and measured the changes made in production of F1 and F2. We predicted that participants would show a conceptually similar compensatory pattern to that observed in Experiment 1: F2 frequency would increase, to offset the perceived acoustical error, but F1 would also change (decrease) to move production changes closer to “hid, bid, did” (see Fig. 1).

Figure 4A shows how F1 and F2 produced by participants, normalized as z-scores, changed during the sensorimotor learning and unlearning phases in response to a 110 mel decrease in F2. By the end of the learning phase, participants had increased their F2 on average by 50 mel to compensate for the decrease in F2 and decreased F1 on average by 14 mel (Figure 4B). Directional t-tests showed that both F1 and F2 frequency at the end of the learning phase (last 30 utterances of learning phase) were significantly different from zero (baseline average) (F2: t(19) = 7.79, p< .001, d = 2.46; F1: t(19) = 2.30, p = .017, d = 0.73).

**Figure 4.**
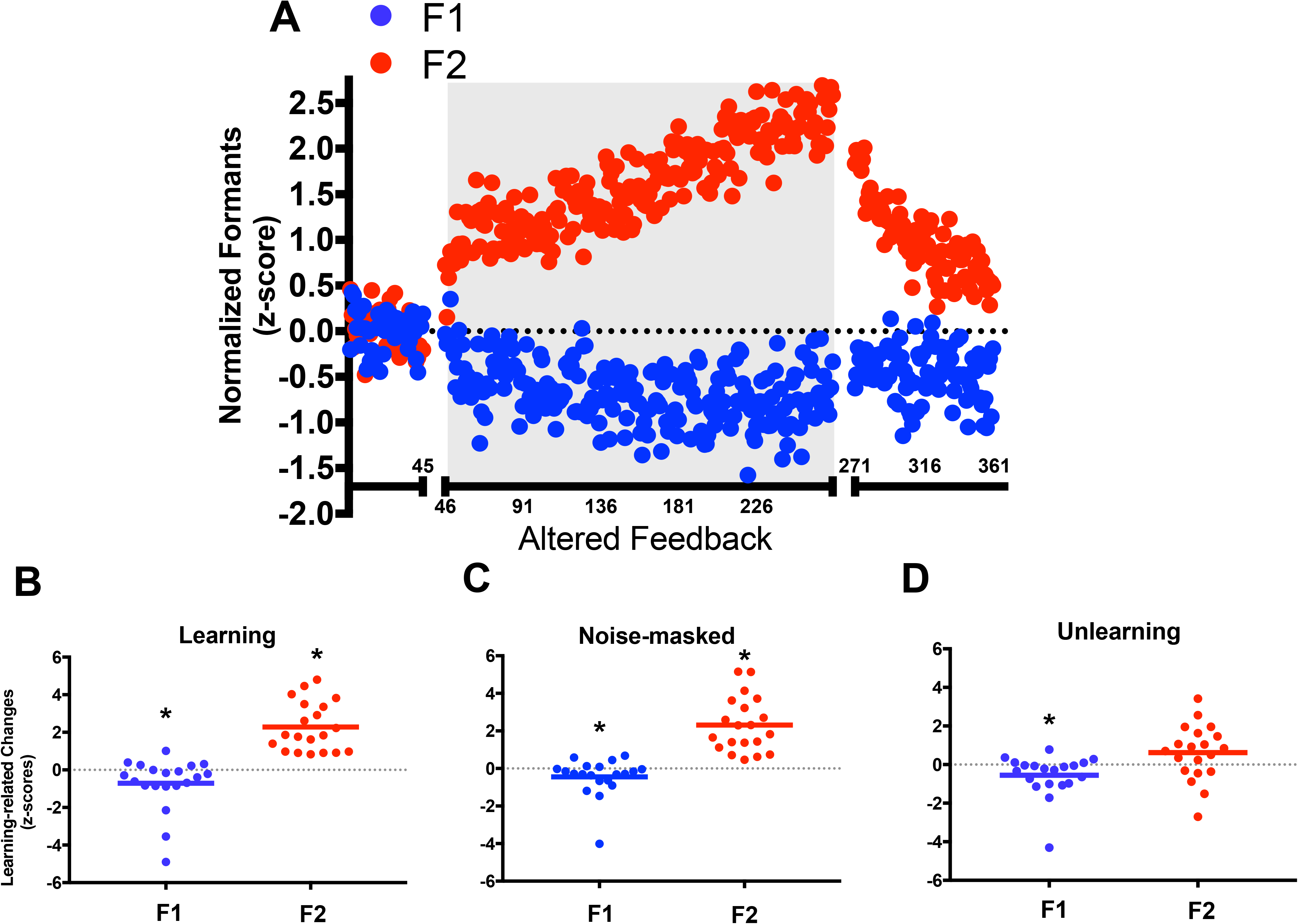
Experiment 2: F2 shifted downwards by 110 mel. See legend to Fig. 3 for details.

**Figure 5.**
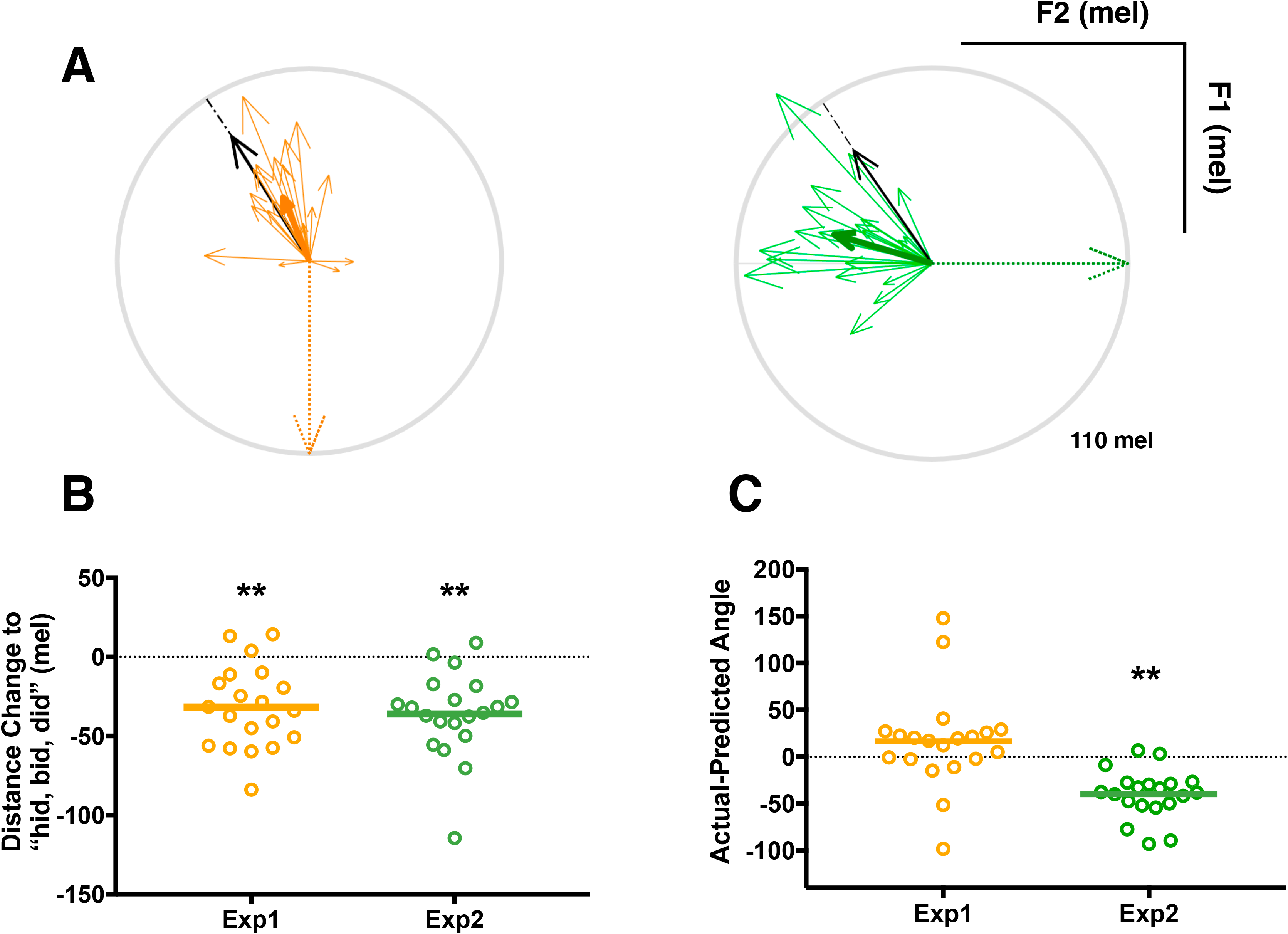
Comparison of changes in vowel production in Experiment 1 and Experiment 2. Top panel: (A) Vector figures of experiment 1 (yellow) and experiment 2 (green). The vectors represent the direction and magnitude of the compensatory change from baseline in vowel space. The thin vectors represent data from individual participants and the thick vectors represent the group average for each experiment. The black arrow represents the average of the direction required to move from “head, bed, dead” to “hid, bid, did”, while the dotted arrow represents the formant frequency shift we applied (F1+110 mel in Experiment 1 and F2−110 mel in Experiment 2). The light grey circle indicates 110 mel. Bottom panel: (B) Learning-related reduction in Euclidean distance between “head, bed, dead” and “hid, bid, did” in F1-F2 space. Negative values indicate that production of “head, bed, dead” by the end of learning has moved closer to “hid, bid, did”. Bottom panel: (C) The difference between the angles of the direction of actual compensatory movement and predicted movement (to “hid, bid, did”) for each participant. Open circles represent individual participants, the solid lines represent the group means. Distance changes (B) and Actual-Predicted angles (C) were compared to zero in each experiment. * p < 0.05 and ** p < 0.01

Figure 4C shows the change in normalized F1 and F2 frequency for heavily noise-masked speech from before and after learning in response to a decrease in F2 feedback. There was a significant increase in F2 production (t(19) = 7.12, p< .001, d = 1.59), and a significant reduction in F1 (t(19) = 1.99, p = .030, d = 0.45) from before to after learning.

By the end of the unlearning phase, F2 frequency returned to baseline (one-sample t-test against baseline average, Z = 0: t (19) = 1.95, p= .066), while F1 frequency remained significantly reduced relative to baseline (t (19) = 2.31, p= .032) after these 90 trials with normal feedback (Fig. 4D).

The results of Experiment 2 provide further support for the idea that feedback perturbations induce changes not only in the shifted formant, but also in the unshifted formant. As in Experiment 1, we evaluated whether these combined changes served to move the produced speech token closer to another vowel sound. The vector analysis confirmed a learning-related reduction in the Euclidean distance between “head, bed, dead” and “hid, bid, did” in F1-F2 space Experiment 2 (Figure 5B). By the end of learning, the productions of “head, bed, dead” were 36 mel closer to the productions of “hid, bid, did” in response to the F2 shift-down (one-sample t-test against zero, t(19) = 5.96, p < .001, d = 1.88). However, the direction of the change was significantly different from the angle predicted (Figure 5C; t(19) = 6.86, p< .001, d = 2.17). Thus, although the results of Experiment 2 met our predictions—a decrease in F2 drove an increase in F2 and a decrease in F1—changes in production were not as well directed towards the “hid, bid, did” part of vowel space as they were in Experiment 1 (Figure 5A). The same results were obtained by analysis of the data from the noise-masked session after learning.

### Comparison of Experiment 1 and 2

We compared the changes in shifted and unshifted formants between the two experiments. The data were rectified as the directions differed between the two experiments (all data were made positive) and analysed using a two-tailed t-test. For the shifted formants, the magnitude of the learning-related change (last 30 utterances of the learning phase) was significantly greater in the F2-shift experiment (Exp 2) compared with the F1-shift experiment (Exp 1) (t(38) = 2.48, p = .018; d = 0.79, Fig. 3B and 4B). However, for the unshifted formants, the magnitudes of the shift at the end of the learning phases of Experiments 1 and 2 did not differ (p = .329; Figs. 3B and 4B). In the vector analysis, there was no significant difference in the magnitude of the learning-related decrease in distance between Experiments 1 and 2 (t(38) = 0.52, p = .610; Fig. 5B). However, the differences between the actual and predicted direction of the changes in each experiment were significantly different (t(38) = 4.385, p < .001; d = 1.39), with a larger difference from the predicted direction for the actual direction of changes in Experiment 2 (Fig. 5C).

Taken together, these findings suggest that the compensatory responses seen in response to shifts applied to F1 and F2 formants in separate experiments each served to move the vowel production towards another vowel in the vowel space and an existing set of articulatory movements associated with that learned production. The significantly larger amount of compensation in the shifted formant (F2) observed in Experiment 2 (in response to a shift down in F2) may explain the discrepancy between the actual and predicted direction required to move from the “head, bed, dead” to the “hid, bid, did” vowel space for that experiment. Nevertheless, the magnitude of the difference between the two vowel productions was significantly reduced following learning in both experiments. It is worth noting that when participants were debriefed after the study, 75% of participants who experienced the F2 shift indicated they were aware of the changes, while only 20% of the participants who experienced the shift upward in F1 noticed them.

## Discussion

The current study investigated how speakers altered their production of the first two formants of vowels by manipulating the auditory feedback for F1 and F2 in two separate experiments. Our hypothesis was that, although the movements required to produce individual formants can be controlled independently to compensate for specific acoustic changes in feedback, speech movements are controlled in a more global way. Since changes in both formants are required to move from one vowel category to another, we predicted that altered auditory feedback of a single formant during speech production would induce changes not only in the shifted formant, but also in the unshifted formant. Furthermore, we predicted that these combined changes would result in a pattern of speech production closer to the frequencies of an existing vowel category.

Consistent with our first prediction, the results of Experiments 1 and 2 showed that by the end of the adaptation phase, participants had significantly changed production of both the shifted and the unshifted formants. Specifically, those who received feedback with an increase in F1, produced speech with a significant decrease in F1 as well as a significant increase in F2. Those who received feedback with a decrease in F2, produced speech with a significant increase in F2 and a significant decrease in F1. The changes relative to speech prior to the application of the shift were evident both in the last 30 trials of the learning phase and in the noise-masked blocks that followed the learning phase (see Figs. 3 & 4).

The results of Experiment 1, in which F1 feedback was increased and participants compensated by decreasing F1 and increasing F2 in their productions, are consistent with previous findings using the same task with a different altered feedback system (Lametti et al., 2018b; Rochet-Capellan and Ostry, 2011). However, the findings of both experiments in the current study contrast with previous work in which participants showed independent changes to the perturbed and unperturbed formants when feedback was gradually shifted in either F1 or F2 (Villacorta et al. 2007). In the latter study, compensatory F1 and F2 changes were observed in the opposite direction to the applied shift in F1 and F2 respectively, as seen in our study. However, F1 did not change during F2 feedback perturbation, which contrasts with the findings for Experiment 2 in our study. Small F2 changes induced by F1 feedback shift were observed by MacDonald et al (2011) but these changes were decreases in F2 when F1 feedback was shifted either up or down. In contrast, in our study, F2 production increased when F1 was shifted up. Furthermore, the small reductions in F2 induced by perturbed F1 in the previous study were found to persist even after the feedback was returned to normal, whereas in our experiments, F2 returned to baseline values but the F1 changes while reduced were not quite returned to baseline.

The difference in the findings from the current study and the previous one by MacDonald and colleagues (2011) may be explained by some experimental differences between the two studies. For example, in the previous study (MacDonald et al., 2011), only 50 utterances were produced during the learning phase, which was assumed to be sufficient for speech adaptation to reach a plateau. In the current study, we witnessed changes in the perturbed and unperturbed formants that continued to occur after 50 utterances (see Figs. 3A and 4A). In our experiments, utterances were of three different words with the same vowel “head, bed, dead” rather than just one word (“head”) used in MacDonald et al., (2011). Intuitively, the varied context should reduce the rate of learning but it could also allow for transfer or generalisation of the adapted vowel to different contexts and possibly a more stable effect by the end of learning. Finally, the magnitude of the shift in F1 feedback was somewhat larger in the MacDonald study (200Hz) than the equivalent of 110 mel, which we applied in Experiment 1 (around 150Hz, but tuned to each participant’s formant to account for the logarithmic relationship between frequency and perception), whereas the F2 shift in the previous study (250Hz) was closer to our Experiment 2 (~265Hz). The degree to which participants were aware of the altered feedback may have affected their compensatory responses in the perturbed formant in particular (though see Munhall et al., 2009). In our Experiment 2, the perturbed formant F2 was noticeably changed (by 75% of participants) and the compensatory response in F2 production was of a much larger magnitude than the change to the perturbed formant F1 in Experiment 1 (only 20% of participants noticed this change). The magnitude of the change in production of the unperturbed formant did not differ between the two experiments, however.

Our findings are compatible with the idea that speech movement parameters can be represented in internal models as coordinated and combined sets to achieve specific sensory goals rather than as precise one-to-one mappings between individual motor dimensions and acoustic outcomes. However, as we discussed above, our results differ from those found previously using a similar paradigm (MacDonald et al., 2011). On the basis of their findings, these authors concluded that the nervous system is capable of independent control of F1 and F2. Even so, the authors suggested that such precise control of single formant may be observed only during feedback error correction and this may not extend to general speech motor control. In our view, the representations in the internal models tested by feedback perturbation paradigms are likely to be the same as those used in general speech motor control. We speculate that the inverse and forward internal models may differ in the extent to which they exert independent and precise articulatory adjustments to each acoustic parameter during feedback perturbation. A previous study showed that stimulation to the cerebellum affected only compensatory responses in F1 production in response to an F1 perturbation, whereas stimulation to the motor cortex affected the amount of compensation in both F1 and F2 in response to the same F1 perturbation (Lametti et al., 2018b). This would suggest that the forward model, which is used to make rapid ongoing adjustments to speech production can independently adjust movements in single dimensions corresponding to separate acoustic changes. On the other hand, the inverse model may encode sensorimotor mappings that represent the combination or combinations of articulatory movements required to achieve a sensory target that exists in the speech repertoire.

Using altered auditory feedback, we showed that adaptive responses to feedback shifts applied to a single formant resulted in changes in production that extended from the shifted to the unshifted formant. These combined changes moved productions closer to the frequencies of nearby existing categories in the vowel space. We conclude that although formants can be controlled independently by adjustments to separate articulatory dimensions, the motor system underlying speech production may prefer to make combined changes in formant production. This latter strategy would serve to maintain speech production within specific regions of the vowel space corresponding to existing speech-sound categories.

## Acknowledgements

This research was supported by a studentship to D-L.T. from the China Scholarship Council. The Wellcome Centre for Integrative Neuroimaging is supported by core funding from the Wellcome Trust (203139/Z/16/Z).

